# SEraster: a rasterization preprocessing framework for scalable spatial omics data analysis

**DOI:** 10.1101/2024.02.01.578436

**Authors:** Gohta Aihara, Kalen Clifton, Mayling Chen, Lyla Atta, Brendan F. Miller, Jean Fan

**Affiliations:** Center for Computational Biology, Whiting School of Engineering, Johns Hopkins University, Baltimore, MD 21211, USA; Department of Biomedical Engineering, Johns Hopkins University, Baltimore, MD 21218, USA

**Keywords:** spatial omics, rasterization, metacells, gene expression, computational biology

## Abstract

**Motivation:** Spatial omics data demand computational analysis but many analysis tools have computational resource requirements that increase with the number of cells analyzed. This presents scalability challenges as researchers use spatial omics technologies to profile millions of cells.

**Results:** To enhance the scalability of spatial omics data analysis, we developed a rasterization preprocessing framework called SEraster that aggregates cellular information into spatial pixels. We apply SEraster to both real and simulated spatial omics data prior to spatial variable gene expression analysis to demonstrate that such preprocessing can reduce resource requirements while maintaining high performance. We further integrate SEraster with existing analysis tools to characterize cell-type spatial cooccurrence. Finally, we apply SEraster to enable analysis of a mouse pup spatial omics dataset with over a million cells to identify tissue-level and cell-type-specific spatially variable genes as well as cooccurring cell-types that recapitulate expected organ structures.

**Availability and implementation:** Source code is available on GitHub (https://github.com/JEFworks-Lab/SEraster) with additional tutorials at https://JEF.works/SEraster.

## 1 Introduction

Spatial omics technologies enable high-throughput molecular profiling of single cells or small groups of cells while preserving their spatial relationships within tissue sections (Bressan et al., 2023). This high-throughput profiling demands computational analysis to leverage both molecular and spatial information in extracting relevant biological insights. Various computational tools have been developed for such analysis, ranging from those that identify spatially variable genes (SVGs) (Kats et al., 2021; Miller et al., 2021; Sun et al., 2020; Svensson et al., 2018; Weber et al., 2022; Zhu et al., 2021) to those that delineate spatial organization and interactions between different cell-types (Cang et al., 2023; Kim et al., 2023; Li et al., 2023; Peixoto et al., 2023; Shao et al., 2022). However, many of these computational tools have runtime and memory requirements that increase with the number of single cells or spatial points analyzed, presenting challenges as technologies continue to improve and researcher apply them to generate large-scale spatial omics data with millions of spatial points. Therefore, preprocessing to streamline such large-scale spatial omics data analyses are needed.

A similar scalability challenge previously emerged for the analysis of single-cell RNA sequencing (scRNA-seq) datasets. To address this problem, preprocessing frameworks were developed to subsample cells while maintaining representative transcriptional heterogeneity (Hie et al., 2019; Ren et al., 2019) or aggregate transcriptionally similar cells into metacells prior to downstream analysis (Baran et al., 2019; Bilous et al., 2022). Both preprocessing techniques by either subsampling or aggregating reduce the number of cells analyzed, thereby lessening the computational resource requirements of downstream analysis and enhancing scalability.

Here, to enhance the scalability of spatial omics data analysis, we developed a preprocessing framework called SEraster to aggregate spatially proximal cells into pixels using rasterization prior to downstream analysis. SEraster further implements sparse matrix representations and parallel processing for enhanced efficiency. We benchmarked the performance of SEraster on downstream spatial omics data analyses including identifying SVGs and cell-type cooccurrence to demonstrate that SEraster enables scalable and accurate analysis of large-scale spatial omics datasets through integration with existing spatial omics data analysis tools. SEraster is implemented as an R package (available on GitHub https://github.com/JEFworks-Lab/SEraster) and uses the SpatialExperiment infrastructure for storing spatial omics data, allowing for streamlined integration with existing spatial omics analysis tools within the R/Bioconductor framework (Righelli et al., 2022).

## 2 Materials and methods

SEraster reduces the number of spatial points in spatial omics datasets for downstream analysis through a process of rasterization where single cells’ gene expression or cell-type labels are aggregated into equally sized pixels based on a user-defined resolution (Figure 1A, Supplementary Information S1). Here, we refer to a particular resolution of rasterization by the side length of the pixel such that finer resolution indicates smaller pixel size and vice versa (Figure 1B). To create a rasterized representation, SEraster initially employs the sf package (Pebesma et al., 2018) to generate pixels by defining square grids that span the x and y spatial coordinate values in the spatial omics dataset. For continuous variables such as gene expression or other molecular information, SEraster aggregates the observed raw counts or normalized expression values for each molecule within each pixel. Such rasterization can also be performed in a cell-type-specific manner by restricting to cells of a particular cell-type prior to rasterization. Alternatively, to create a rasterized representation of categorical variables such as cell-type or cluster labels, SEraster first converts the labels to a model matrix using a one-hot encoding and then treats the model matrix as a features-by-observations matrix to aggregate the number of cells for each label within each pixel. In both applications, aggregation function can be chosen from mean or sum to accommodate user-defined purposes. This rasterization process is implemented in a pixel-wise manner, which can be parallelized with the BiocParallel package (Morgan et al., 2023). Compared to other R packages that can perform rasterization on vector data represented as dense matrices, such as terra (Hijmans et al., n.d.) or stars (Pebesma et al., n.d.), SEraster can rasterize spatial omics datasets represented as either dense or sparse matrices with the Matrix package (Bates et al., n.d.). Since features-by-observations matrix and model matrix are often sparse, this feature further allows SEraster to reduce resource requirements upon rasterization preprocessing. In addition, since rasterized values may be sensitive to edge effects such as the specific boundaries of grids upon rasterization, SEraster enables permutation by rotating the dataset at various angles before rasterization (Figure 1C, Supplementary Information S2). The rasterized output is returned as a SpatialExperiment object, allowing for streamlined integration with existing spatial omics analysis tools within the R/Bioconductor framework (Righelli et al., 2022) for downstream analyses.

**Figure 1.**
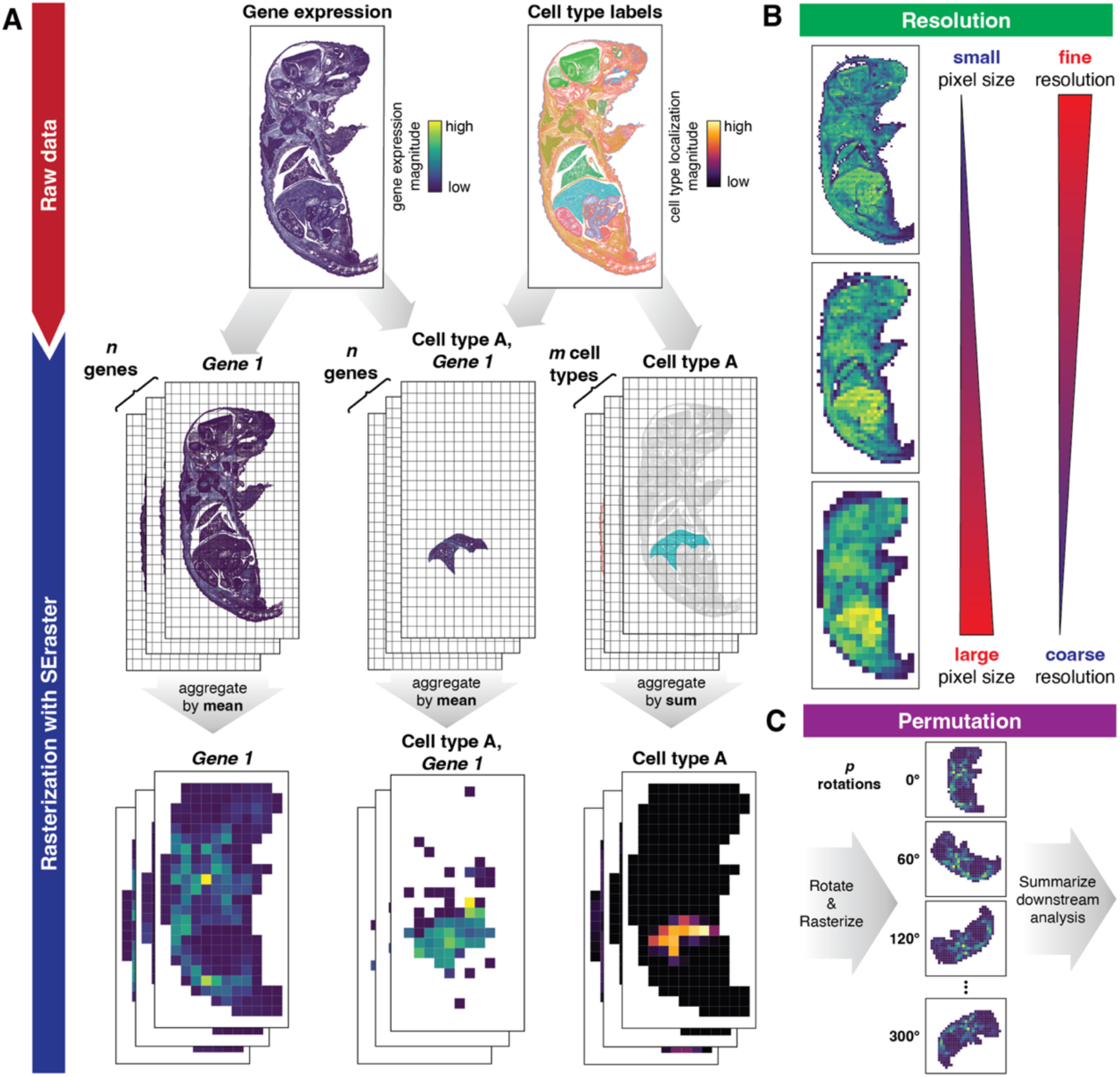
Overview of SEraster. **A.** SEraster reduces the number of spatial points in a given spatial omics dataset prior to downstream analysis by rasterizing or aggregating single cells’ gene expression or cell-type labels into equally sized pixels. SEraster can be applied to aggregate gene expression in a label-specific (e.g. cell-type or cluster) manner as well. **B.** SEraster allows users to control the rasterization resolution or the side length of the pixel. Finer resolution corresponds to a smaller pixel size and coarser resolution corresponds to a larger pixel size. **C.** SEraster enables permutation by rotating the dataset at various angles before rasterization. Downstream analyses performed on rotated datasets can be summarized to help control for edge effects.

To explore the potential utility of rasterization in spatial omics analysis, we apply SEraster as a preprocessing step prior to downstream spatial omics analysis using both simulated and real spatial omics data. In particular, we benchmark the impact of rasterization on runtime and accuracy in identifying SVGs with the nnSVG package (Weber et al., 2022). We further demonstrate how rasterized cell-types can be used with the CooccurrenceAffinity package (Mainali et al., 2022; Mainali & Slud, 2022) to recapitulate expected pairs of cell-types that tend to spatially cooccur. In this manner, SEraster can be used as a pre-processing step to enable scalable and accurate analysis of large-scale spatial omics datasets with existing tools.

## 3 Results

### 3.1 Rasterization reduces runtime while maintaining accuracy in the identification of spatially variable genes

To evaluate the potential utility and impact of rasterization on the performance of downstream analysis, we focused on identifying SVGs within tissues. A number of computational tools have been previously developed to identify SVGs (Kats et al., 2021; Miller et al., 2021; Sun et al., 2020; Svensson et al., 2018; Weber et al., 2022; Zhu et al., 2021). We applied SEraster and one of these methods, nnSVG, to a single-cell resolution spatial transcriptomics dataset of a coronal section of the mouse brain containing 83,546 cells assayed by MERFISH (Figure 2A, Supplementary Information S3i, (Vizgen, n.d.)). We first evaluated the runtime of SVG analysis without parallelization when applied to the dataset at single-cell resolution (sc) versus rasterized at 50 µm, 100 µm, 200 µm, and 400 µm resolutions (Figure 2B). Combining SEraster and nnSVG reduced the total runtime to 26.8%, 9.9%, 3.9%, and 2.7% of that when running nnSVG at single-cell resolution for 50 µm, 100 µm, 200 µm, and 400 µm rasterization resolutions, respectively (Figure 2C, Supplementary Information S4). This shorter runtime is expected due to the fewer numbers of spatial points considered, particularly at coarser resolutions with larger pixel sizes (Supplementary Figure 1A). We also characterized the performance of nnSVG in identifying SVGs when applied to rasterized gene expression by comparing SVGs identified at single-cell resolution, which were treated as ground truth, to those detected at each rasterized resolution (Supplementary Information S4). We find that true positive rate (TPR) and positive predictive value (PPV)–also known as sensitivity and precision, respectively–remained high across rasterization resolutions, ranging from 0.96 to 0.99 for TPR and from 0.92 to 0.96 for PPV (Figure 2D). True negative rate (TNR)–also known as specificity–ranged from 0.60 to 0.80 across resolutions. Consistently, we observed high correlation between the ranks of each gene based on nnSVG’s likelihood statistics between single-cell and each rasterized resolution (Figure 2E). nnSVG’s gene rankings indicate the strength of the spatial gene expression patterns. Hence, the high correlation of gene rankings suggests that the spatial pattern strengths of genes are generally retained even with SEraster preprocessing. We observe lower correlations at coarser resolutions, suggesting that the forementioned relationships were less well retained at coarser resolutions. These results suggest that rasterization can generally preserve high performance while decreasing runtime for SVG analysis.

**Figure 2.**
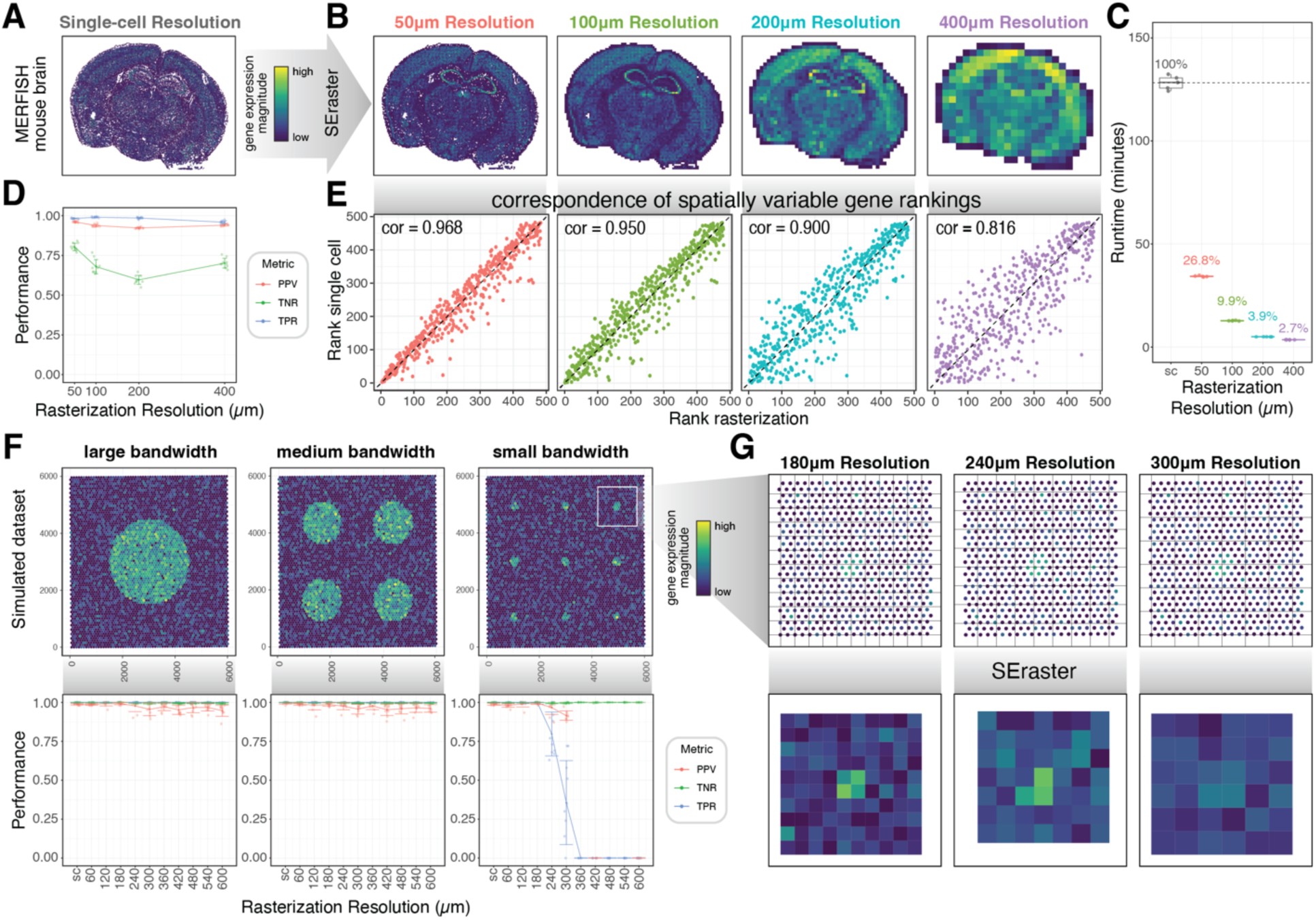
Impact of rasterization on the identification of spatially variable genes (SVGs). **A.** Spatial transcriptomics dataset of a coronal section of mouse brain assayed by MERFISH shown at single-cell resolution colored by log-normalized total gene expression per cell. **B.** MERFISH mouse brain dataset rasterized at 50 µm, 100 µm, 200 µm, and 400 µm resolutions colored by sum of rasterized gene expression per pixel. **C.** Time (in minutes) required to run nnSVG at single-cell (sc) resolution and SEraster preprocessing with nnSVG at selected rasterized resolutions (𝑛 = 5 for each resolution). Boxplots represent medians, first, and third quartiles, and whiskers extend to values no further than 1.5 times the interquartile range from each quartile. Percentage values of runtime at rasterized resolutions compared to that at single-cell resolution are shown. **D.** Performance in terms of Positive Predictive Value (PPV), True Negative Rate (TNR), and True Positive Rate (TPR) evaluated by comparing predicted SVG and non-SVGs at single-cell versus selected rasterization resolutions for the MERFISH mouse brain dataset. Line plots show mean and standard deviation across 10 permutations. Data points are indicated by dots. **E.** Correspondence of gene rankings based on the estimated LR statistic from nnSVG at single-cell resolution and selected rasterized resolutions. Corresponding Spearman’s correlation coefficients are shown as text labels. **F.** Simulated SVG dataset shown at single-cell resolution with large, medium, and small bandwidths corresponding to circular spatial patterns for SVGs with radii of 1500 µm, 750 µm, and 150 µm (top). Each single-cell is colored by log normalized gene expression of ground truth SVG. Performance metrics in terms of PPV, TNR, and TPR are computed by comparing ground truth versus predicted SVG and noise SVG labels for each gene at single-cell (sc) and rasterized resolutions ranging from 60 to 600 µm (bottom). Line plots show mean and standard deviation across 10 permutations. Data points are indicated by dots, and NaN values are removed. **G.** Close-up visualizations of SVG with small bandwidth at single-cell resolution colored by log normalized gene expression and grid used upon rasterization at selected resolutions (top). Close-up visualizations of rasterized gene expression for the same gene after rasterization at selected resolutions (bottom).

To further characterize the effects of rasterization resolution on nnSVG’s performance, we simulated spatial omics data with 100 SVGs and 900 noise genes across 4992 spatial points using a previously developed simulation framework (Supplementary Information S5i, (Weber et al., 2022)). By using simulations, we were able to modulate the scale of spatial patterns (large, medium, and small corresponding to circular spatial patterns for SVGs with radii of 1500 µm, 750 µm, and 150 µm respectively) (Figure 2F). We evaluated performance by comparing the simulated ground truth SVG or noise labels for each gene to those predicted by nnSVG when applied at single-cell versus at rasterized resolutions ranging from 60 to 600 µm (Supplementary Information S4). For simulated datasets with large and medium spatial patterns, nnSVG’s performance with rasterization remained high, with TPR values consistently at 1, PPV ranging from 0.95 to 1, and TNR ranging from 0.99 to 1 across evaluated resolutions (Figure 2F). Notably, at all evaluated resolutions, the rasterized pixel size was smaller than the simulated SVGs’ circular spatial patterns with radii of 1500 µm and 750 µm. However, for simulated datasets with small spatial patterns, TPR started decreasing at 240 µm resolution and reached 0 at 360 µm resolution (Figure 2F) due to nnSVG failing to detect any SVG at 300 µm and coarser resolutions, resulting in undefined PPV values and high TNR. These results are expected since the simulated SVGs’ circular spatial pattern has a radius of 150 µm. At coarser resolutions, the rasterized pixel size is too large such that cells with high expression of SVGs are aggregated with those with low expression of SVGs, eliminating signals (Figure 2G). These findings suggest that users should choose a rasterization resolution that is sufficient to capture the size of spatial patterns of interest in order to maintain accuracy in their SVG analysis when using rasterization as a preprocessing step.

### 3.2 Rasterization enables scalable characterization of spatial cell-type cooccurrence

To demonstrate additional potential applications of rasterization, we sought to use SEraster in identifying spatially cooccurring cell-types. We applied SEraster to a single-cell resolution spatial proteomics dataset of a human spleen containing 154,446 cells assayed by CODEX (Figure 3A, Supplementary Information S3ii, (Currlin et al., 2022)). We used SEraster to aggregate cell counts for each cell-type within pixels at 100 µm resolution. We then treated rasterized cell-type counts in a classic framework of balls (cell-types) placed in boxes (100 µm pixels) to characterize cell-type cooccurrence (Supplementary Information S6). Briefly, rasterized cell-type counts were used to compute a relative enrichment (RE) metric, or the ratio of observed to expected cell-type counts, per pixel to account for variability in cell density and cell-type proportions (Figure 3B). Each pixel’s RE value was binarized based on a selected threshold (Figure 3B). Based on the binarized data, we used the CooccurrenceAffinity package to compute the maximum likelihood estimate of the affinity metric, 𝛼̂, for each cell-type pair, with positive 𝛼̂ indicating cooccurrence and negative 𝛼̂ indicating separation (Mainali et al., 2022; Mainali & Slud, 2022). We identified cell-type pairs with statistically significant coocurrence (𝛼̂ > 0, p-value ≤ 0.05) (Figure 3C). Hierarchical clustering of 𝛼̂ values demonstrated that this approach can recapitulate expected spatial niches such as the white pulp and red pulp, previously identified by other analysis approaches (Peixoto et al., 2023) as well as visually validated at single-cell resolution (Figure 3D, Supplementary Information S6).

**Figure 3.**
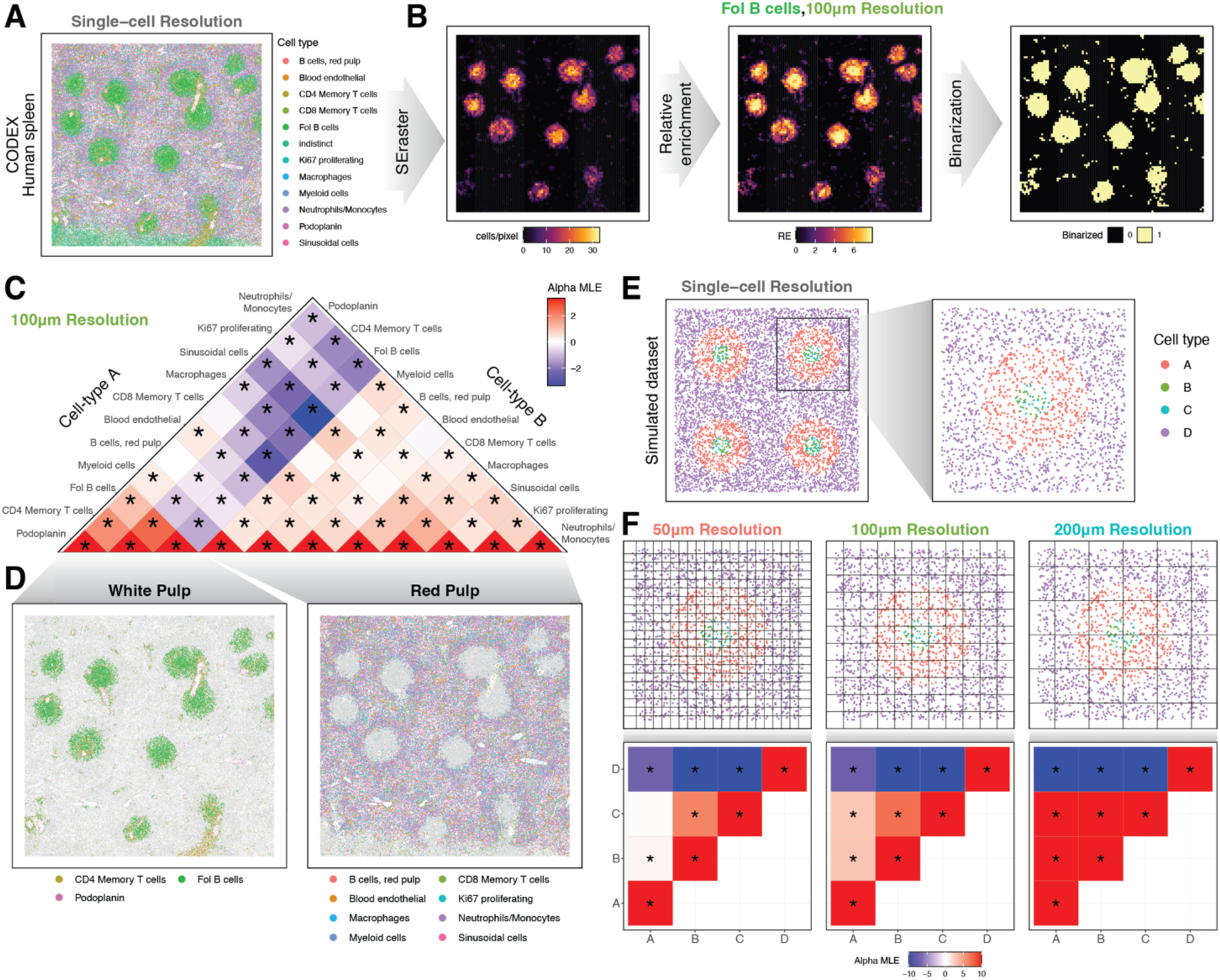
Rasterization of cell-type labels enables analysis of cell-type cooccurrence. **A.** Spatial proteomics dataset of the human spleen assayed by CODEX shown at single-cell resolution colored by cell-types. **B.** Follicular B cells in the CODEX human spleen dataset shown as rasterized cell-type count, relative enrichment metric, and binarized value per pixel at 100 µm resolution (from left to right). **C.** Summary of cell-type cooccurrence analysis with SEraster and CooccurrenceAffinity. Heatmap is colored by the maximum likelihood estimate of the affinity metric (alpha MLE or 𝛼̂) for corresponding cell-type pairs. Maximum and minimum 𝛼̂ values are winsorized at max (|𝛼̂|) of non-self-cell-type pair for better visualization. Statistically significant cooccurrences or separations (p-value ≤ 0.05) are indicated by asterisks (*). **D.** Spatial niches identified from the cell-type cooccurrence analysis are visualized at single-cell resolution colored by cell-types. Based on cell-type compositions, two niches are labeled as white pulp (left) and red pulp (right). **E.** Simulated cooccurrence dataset shown at single-cell resolution colored by cell-types. **F.** Close-up visualizations of cell-type cooccurrence patterns at single-cell resolution colored by cell-types with the grid used upon rasterization shown at selected resolutions (top). Summarized results of cell-type cooccurrence analysis with SEraster and CooccurrenceAffinity at selected rasterization resolutions. Heatmap is colored by 𝛼̂ , and statistically significant cooccurrences or separations are indicated by asterisks (*) (bottom).

To further evaluate the impact of rasterization resolution on cell-type cooccurrence analysis, we employed a previously developed simulated dataset that mimics cell-type localizations at various scales (Peixoto et al., 2023). In this dataset, cell-types B and C are spatially intermixed in circular spatial patterns with radius of 100 µm, and cell-type A surrounds cell-types B and C in doughnut-shaped spatial patterns with radii ranging from 100 µm to 300 µm, separating them from cell-type D (Figure 3E, Supplementary Information S5ii). At a rasterization resolution of 50 µm, cell-types B and C had high, positive 𝛼̂ while A and B as well as A and C had 𝛼̂ ≈ 0 (Figure 3F). This is expected because 50 µm resolution is smaller than the size of cell-type A’s doughnut-shaped structures. As a result, pixels did not capture the spatial pattern that cell-type A surrounds cell-types B and C. With coarser resolutions—for instance, 100 µm and 200 µm resolutions that are within the range of cell-type A’s doughnut-shaped structures—pixel size is large enough to capture spatial patterns formed by cell-types A, B, and C in one pixel. Thus, cell-types A and B as well as A and C were identified as cooccurring (Figure 3F). These results suggest that changing rasterization resolutions can capture cell-type cooccurrence relationships at various spatial length scales. Overall, SEraster can transform spatial omics data with cell-type labels into a classic balls-in-boxes formulation to enable characterization of cell-type cooccurrence.

### 3.3 Rasterization enables spatial analysis of spatial transcriptomics data of a whole mouse pup with over a million cells

Having demonstrated that our approach works as expected, we applied SEraster to a single-cell resolution spatial transcriptomics dataset of a whole mouse pup containing 1,330,087 cells assayed by Xenium (Figure 4A, Supplementary Information S3iii, (10X Genomics, 2023)). To identify SVGs, we again attempted to use nnSVG. However, running nnSVG, which scales linearly with the number of spatial points, at single-cell resolution failed to complete within 24 hours (Supplementary Information S4). On the other hand, rasterizing the dataset to 100 µm resolution using SEraster and running nnSVG required an average total runtime of 54 ± 4 minutes without parallelization (𝑛 = 5, error is computed as standard deviation, Supplementary Information S4), further highlighting the potential utility of SEraster in reducing computational resource requirements. We thus performed SVG analysis on the entire tissue rasterized at 100 µm resolution. Expectedly, all 379 profiled genes were identified as SVGs given that these genes were chosen to identify specific organs and tissue regions, which are highly spatially compartmentalized.

**Figure 4.**
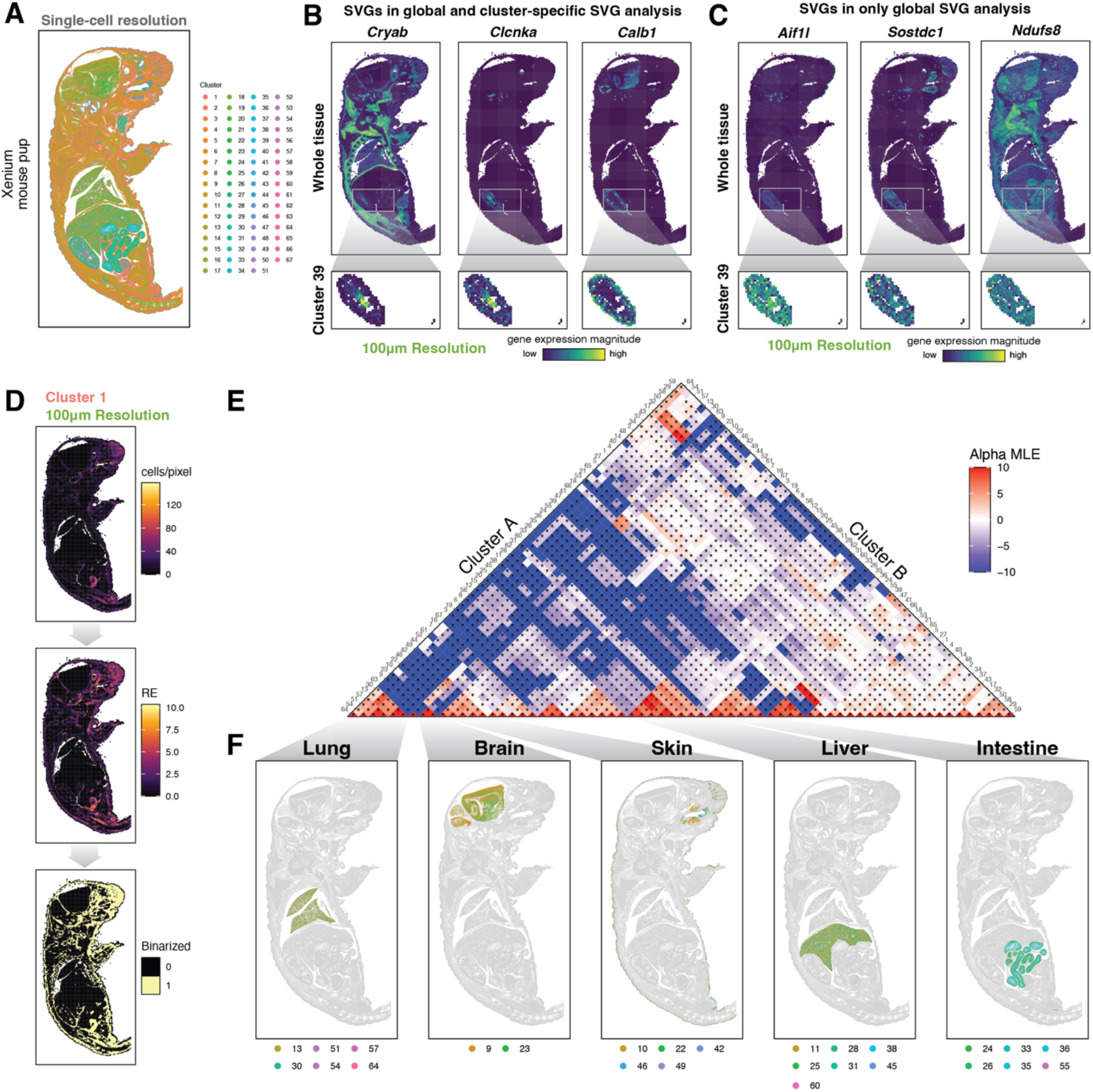
Spatial analysis of a whole mouse pup with over a million cells. **A.** Spatial transcriptomics data of a whole mouse pup assayed by Xenium shown at single-cell resolution colored by clusters from graph-based clustering. **B.** Rasterized gene expression of *Cryab* (left), *Clcnka* (middle), *Calb1* (right) at 100 µm resolution for the whole tissue and within cluster 39. **C.** Rasterized gene expression of *Aifl1* (left), *Sostdc1* (middle), *Ndufs8* (right) at 100 µm resolution for the whole tissue and within cluster 39. **D.** Cluster 1 in the Xenium whole mouse pup dataset shown as rasterized cluster count, relative enrichment metric, and binarized value per pixel at 100 µm resolution (from top to bottom). **E.** Summary of cluster cooccurrence analysis with SEraster and CooccurrenceAffinity. Heatmap is colored by the maximum likelihood estimate of the affinity metric (alpha MLE or 𝛼̂) for corresponding cluster pairs. Statistically significant cooccurrences or separations (p-value ≤ 0.05) are indicated by asterisks (*). **F.** Spatial niches identified from the cluster cooccurrence analysis are visualized at single-cell resolution colored by clusters. Based on the spatial locations of cooccurring clusters, these spatial niches are labeled as lung, brain, skin, liver, and intestine (left to right).

To better understand spatial gene expression variation within specific organs and tissue regions, we performed cluster-specific SVG analysis at 100 µm resolution for each transcriptionally distinct cell-cluster previously identified through graph-based transcriptional clustering (Supplementary Information S4, (10X Genomics, n.d.)). As an example, we focused on Cluster 39, which putatively corresponds to the kidney based on its spatial location and differentially upregulated genes (Supplementary Information S3iii). Rasterizing just cells corresponding to cluster 39, we again performed SVG analysis to identify 118 SVGs. Among these SVGs included *Cryab*, *Clcnka*, *Calb1*, which exhibited statistically significant spatial variation (p-value ≤ 0.05) both in the whole tissue analysis and cluster-specific analysis (Figure 4B). On the other hand, genes such as *Aif1l*, *Sostdc1*, *Ndufs*8 only exhibited statistically significant spatial variation (p-value ≤ 0.05) in the whole tissue analysis and not in the cluster-specific analysis as these genes exhibit more uniform expression within the cluster (Figure 4C). These results demonstrate that SEraster can be used to help identify SVGs in the whole tissue as well as in a cluster-specific manner.

We further applied SEraster with CooccurrenceAffinity to characterize cell-cluster cooccurrence and identify spatial niches in the whole mouse pup at 100 µm resolution (Figure 4D, Supplementary Information S6). We evaluated all 2278 possible cell-cluster pairs to identify 475 with statistically significant cooccurrence (𝛼̂ > 0, p-value ≤ 0.05) in 10 ± 1 minutes without parallelization (𝑛 = 5, error is computed as standard deviation, Supplementary Information S4), again underscoring the scalability of our rasterization-based framework (Figure 4). We further performed hierarchical clustering of 𝛼̂ values to find groups of cell-clusters that cooccur visually correspond to spatially distinct organ structures (Figure 3F, Supplementary Information S6). Further, we observed that such cell-cluster cooccurrence patterns forming spatially distinct organ structures are robust across rasterization resolutions ranging from 50 µm to 400 µm resolutions (Supplementary Figure 2), demonstrating the stability of these cell-cluster cooccurrence relationships.

These results demonstrate that rasterization preprocessing with SEraster can be applied to large-scale spatial omics datasets with over a million single cells to enable the identification of SVGs at the whole tissue as well as cluster-specific levels and detection of cell-cluster cooccurrence patterns that correspond to spatially distinct organ structures.

## 4 Discussion

Analysis of spatial omics data provide researchers with means to delineate spatial patterns of molecular and cellular organizations. To improve the scalability of spatial omics data analysis, we developed SEraster to use rasterization as a preprocessing step to reduce the number of spatial points prior to downstream analysis by aggregating continuous variables, such as gene expression, or categorical variables, such as cell-type labels, at single-cell resolution into equally sized pixels based on a user-defined resolution. Such reduction in the number of spatial points enabled the spatial analysis of a whole mouse pup spatial omics dataset with over a million single cells using existing tools that would not have otherwise been computationally tractable. Applying SEraster prior to SVG analysis with nnSVG, we find that SEraster reduces runtime requirements without significantly compromising performance compared to single-cell resolution. Likewise, SEraster also enabled rapid, pair-wise cell-type colocalization with CooccurrenceAffinity.

While we exclusively examined single-cell resolution imaging-based spatial omics datasets in this paper, the same framework can, in principle, be applied to other non-single-cell resolution spatial omics data (Bressan et al., 2023; Moffitt et al., 2022), though interpretation of results will need to consider the lack of single-cell resolution and potential confounding due to cell-type mixtures. Likewise, although we have focused on applying SEraster to spatial transcriptomics and proteomics data, the framework can be applied to other spatial omics technologies as well. Further, while we have focused on applying rasterization on individual spatial omics datasets, SEraster can also be applied to multiple tissue sections in order to create shared pixels in the same coordinate framework (Supplementary Information S1). For instance, such analysis facilitates spatial molecular comparisons after structural alignment (Supplementary Figure 3, (Clifton et al., 2023)).

Several limitations of SEraster should be considered when integrating SEraster with downstream analysis tools. As shown with SVG analysis on simulated datasets, rasterization resolution is a user inputted hyperparameter that may affect downstream analysis. A rasterization resolution that is too coarse to capture the size of spatial patterns of interest will result in false negatives. Here, we draw a parallel between rasterization resolution and the concept of ‘grain size’ in ecology (Wiens, 1989), the spatial scale at which ecological processes are studied. As the choice of grain size will depend on the ecological phenomena of interest, so will the choice of rasterization resolution depend on the biological processes of interest. We recommend optimizing rasterization resolution based on prior knowledge regarding the scale of biological processes being studied.

Although we have focused on demonstrating the utility of rasterization preprocessing with SEraster in downstream spatial omics analyses such as identifying SVGs and pair-wise cell-type cooccurrence, rasterization may also be amenable as a preprocessing step for other types of spatial omics analysis. For example, a few computational tools for inferring cell-cell communication have been developed for spatial transcriptomics data (Cang et al., 2023; Kim et al., 2023; Li et al., 2023; Shao et al., 2022). Potential future work may involve integrating these methods that leverage distributions of molecular information with SEraster to enable more scalable cell-cell communication analysis. However, we caution that additional statistical characterization is needed to benchmark performance and mitigate false positives.

Finally, as spatial omics datasets continue to increase in size, in the future, we anticipate spatial omics datasets may need to be stored in standardized data infrastructure with lazy representation of larger-than-memory data such as the Zarr file format used in the SpatialData Python library (Marconato et al., 2023). SEraster can potentially be integrated with such data infrastructure to enable rasterization of larger-than-memory data in spatially-indexed chunks rather than loading the entire dataset into memory.

To conclude, SEraster improves scalability by reducing the number of spatial points and broadens statistical methods for spatial omics data explorations. As spatial omics datasets continue to increase in the number of spatial points assayed, integrating such rasterization preprocessing with existing and new computational tools may enable more efficient analysis.

## Acknowledgements

This work was supported by the National Science Foundation under Grant No. 2047611 and by the HuBMAP Integration, Visualization & Engagement (HIVE) Initiative under Award Number 1OT2OD026673.

## Supplementary Information for

### Supplementary Methods

#### 1. SEraster Overview

For both continuous variables, such as gene expression, and categorical variables, such as cell-type labels, the process of rasterization only differs by aggregation function. For a given spatial omics dataset with a features-by-observations matrix and x, y spatial coordinates, SEraster creates square grids with side length of each pixel corresponding to user-defined resolution. The sf package (Pebesma et al., 2018) is used to create these square grids based on the bounding box including all spatial points in a given dataset as well as the surrounding space. Size of the surrounding space is approximately half of the user-defined resolution to ensure that all cells would be incorporated into square grids. SEraster’s rasterization is implemented in a pixel-wise manner, which is parallelized with the BiocParallel package. For each pixel with at least one cell, the features-by-observations matrix is subset to cells that reside within the pixel of interest, and their data is aggregated by mean or sum, essentially creating a features vector for the pixel of interest. At the end, a new features-by-observations matrix as well as x,y spatial coordinates of pixel centroids for rasterized dataset are returned as a SpatialExperiment object with the SpatialExperiment package.

Since both gene counts matrix and model matrix are sparse, they are often represented as sparse matrices to reduce memory requirements. Hence, SEraster can rasterize a SpatialExperiment object with the features-by-observations matrix represented as either dense or sparse matrix (dgCMatrix) using the Matrix package.

Further, SEraster can simultaneously rasterize multiple spatial omics datasets with shared pixel coordinates. To do so, SEraster creates square grids based on a common bounding box defined by minimum and maximum x,y spatial coordinates across given spatial omics datasets.

#### 2. Permutation

To address the impact of specific locations and orientations of grids on downstream analysis, SEraster conducts permutation by rotating the single-cell resolution dataset at various angles, 𝜃, prior to rasterization with the following rotation matrix.

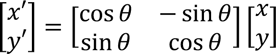

where 𝜃 is determined by a counterclockwise rotation from the x-axis around the midpoint of a two-dimensional Cartesian coordinate system.

#### 3. Biological datasets

##### i. MERFISH mouse brain dataset

The MERFISH dataset of 1 mouse brain coronal section (slice 2, replicate 1) was obtained from the Vizgen website for *MERFISH Mouse Brain Receptor Map data release* (Vizgen, n.d.). “Blank” genes used for quality control were removed from the dataset. If cells did not have any gene with at least 2 RNA counts, they were considered as poor quality cells and were removed from the dataset. These filtering steps resulted in 83,546 cells and 483 genes. RNA counts were normalized by cell volume (divided per cell by the corresponding cell volume and scaled by the mean of all cell volume) and log_10_ transformed with a pseudocount of 1. Raw RNA counts, processed gene expression, and spatial coordinates were used to construct a SpatialExperiment object.

##### ii. CODEX human spleen dataset

The pre-processed CODEX dataset containing 154,446 cells and 29 protein markers for 1 human spleen tissue section was obtained from the HuBMAP Data Portal corresponding to dataset IDs: HBM389.PKHL.936 (Currlin et al., 2022). Protein expression was normalized by cell area and log_10_ transformed with a pseudocount of 1. Cell-type labels for the same dataset were obtained from the CRAWDAD package (Peixoto et al., 2023). Processed protein expression, cell-type labels, and spatial coordinates were used to construct a SpatialExperiment object.

##### iii. 10X Genomics Xenium whole mouse pup dataset

The Xenium dataset of 1 day old mouse pup section was obtained from the 10X Genomics website for *Whole Mouse Pup Preview Data* (10X Genomics, 2023). Results for graph-based clustering analyses as well as differential gene expression analysis were also obtained from the same website. Genes used for quality control (those not included in the gene panel) were removed from the dataset. If cells did not have any gene with at least 2 RNA counts, they were considered as poor quality cells and were removed from the dataset. These filtering steps resulted in 1,330,087 cells and 379 genes. RNA counts were normalized by nucleus area (divided per cell by the corresponding nucleus area and scaled by the mean of all nucleus area) and log_10_ transformed with a pseudocount of 1. Raw RNA counts, processed gene expression, cluster labels (graph-based clustering), and spatial coordinates were used to construct a SpatialExperiment object.

Graph-based clustering labels were used for cluster-specific spatial variable gene (SVG) analysis, and an example of cluster-specific SVG analysis was shown using cluster 39. Cluster 39 was determined to likely correspond to kidney due to its location within the whole mouse pup tissue and differentially expressed genes (DEGs). Based on the DEG analysis performed by 10X Genomics (10X Genomics, n.d.), cluster 39 differentially upregulated *Lrp2*, a marker for proximal tubules (Khundmiri et al., 2021).

#### 4. Spatial variable gene (SVG) analysis

SEraster aggregates gene expression of a given spatial omics dataset and creates a new SpatialExperiment object with a new features-by-observations matrix and x,y coordinates of pixel centroids. To conduct SVG analysis after rasterization with SEraster, this SpatialExperiment object is directly used as an input to the nnSVG package (Weber et al., 2022). For the whole tissue SVG analysis, SEraster rasterizes the entire dataset. For cell-type- or cluster-specific SVG analysis, the entire dataset is subsampled to a cell-type or cluster of interest, and SEraster rasterizes the subsampled dataset.

Runtime was measured using Sys.time function in the R Base package and averaged across 5 trials. While single-cell resolution quantified runtime for nnSVG alone, rasterization resolution quantified runtime for SEraster alone, nnSVG alone, and combined. This runtime analysis was done on a Mac Studio Apple M2 Ultra with 192GB unified memory (24-core CPU, 60-core GPU, 32-core Neural Engine) using 1 worker for single computer multicore parallel evaluation (BiocParallel::MulticoreParam(workers = 1)).

Performance metrics were computed by comparing Boolean labels for each gene, indicating whether it is a SVG or not, between ground truth condition and selected rasterized resolutions. Since the nnSVG package computes multiple-testing-p-value using the likelihood ratio (LR) test with 2 degrees of freedom within itself (Weber et al., 2022), genes are considered as SVGs or to have statistically significant spatial variation based on a multiple-testing-p-value cutoff of 0.05 or below. For the MERFISH mouse brain dataset, Boolean labels for the single-cell resolution were used as ground truths and compared with those for the rasterization resolutions.

For the simulated SVG dataset, known Boolean labels were compared with those for the single-cell and rasterization resolutions. The following performance metrics were computed.

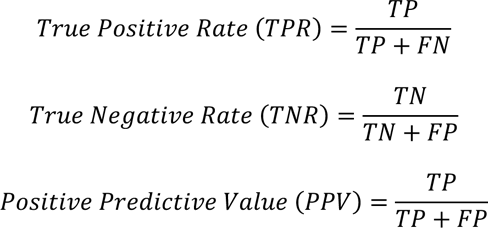

where 𝑇𝑃, 𝑇𝑁, 𝐹𝑃, 𝐹𝑁 denote the number of true positives, true negatives, false positives, and false negatives, respectively. To accommodate the sensitivity of rasterization from the orientation of grids with respect to the spatial omics data, performance metrics were summarized across 10 permutations rotated at 0°, 36°, 72°, 108°, 144°, 180°, 216°, 252°, 288°, and 324°.

Further, for the MERFISH mouse brain dataset (rotated at 0°), gene rankings based on the estimated LR statistics computed within nnSVG were compared between single-cell resolution and selected rasterization resolutions. Spearman’s correlation coefficient was computed for each comparison.

#### 5. Simulated datasets

##### i. nnSVG

Simulated datasets for SVG analysis were obtained from nnSVG (v1.5.8) (Weber et al., 2022). All datasets contained 4992 spatial points. These datasets simulated 100 SVGs and 900 noise genes based on the Visium human DLPFC dataset with 3 varying scales of spatial patterns (Weber et al., 2022). For all datasets, we scaled the spatial coordinates to 6,000 µm and 6,000 µm, which resulted in the scale of circular spatial patterns to have radii of 1500 µm, 750 µm, and 150 µm.

##### ii. CRAWDAD

The simulated dataset for cell-type cooccurrence analysis was obtained from CRAWDAD (v1.0) (Peixoto et al., 2023). This simulated dataset contains 8,000 cells that are randomly distributed across 2,000 µm x 2,000 µm space.

#### 6. Cell-type cooccurrence analysis

SEraster uses a two-step binarization approach to create binary presence/absence data for a given spatial omics dataset. Relative enrichment (RE) metric represents the ratio of observed to expected cell-type counts, and similar framework has been used in the context of single-molecule localization microscopy for assessing colocalization of two molecular species with Voronoï tessellation (Ejdrup et al., 2022). In SEraster, RE for each pixel, 𝑖, and category, 𝑗, i computed using the rasterized cell-type or cluster labels based on the following equation

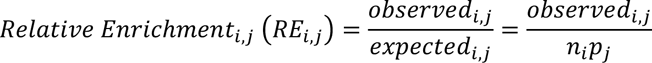

where 𝑛_1_ denotes the total number of cells in pixel 𝑖 and 𝑝_̂_ denotes the proportion of category 𝑗. Relative enrichment metric is then binarized based on a threshold of 1.

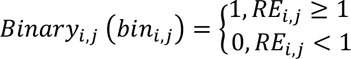

This approach accommodates the differences in the proportion of cell-types and the cell density variance across spatial coordinates.

Binarized dataset is used to create 2 x 2 contingency tables indicating the presence and absence of corresponding cell-types.

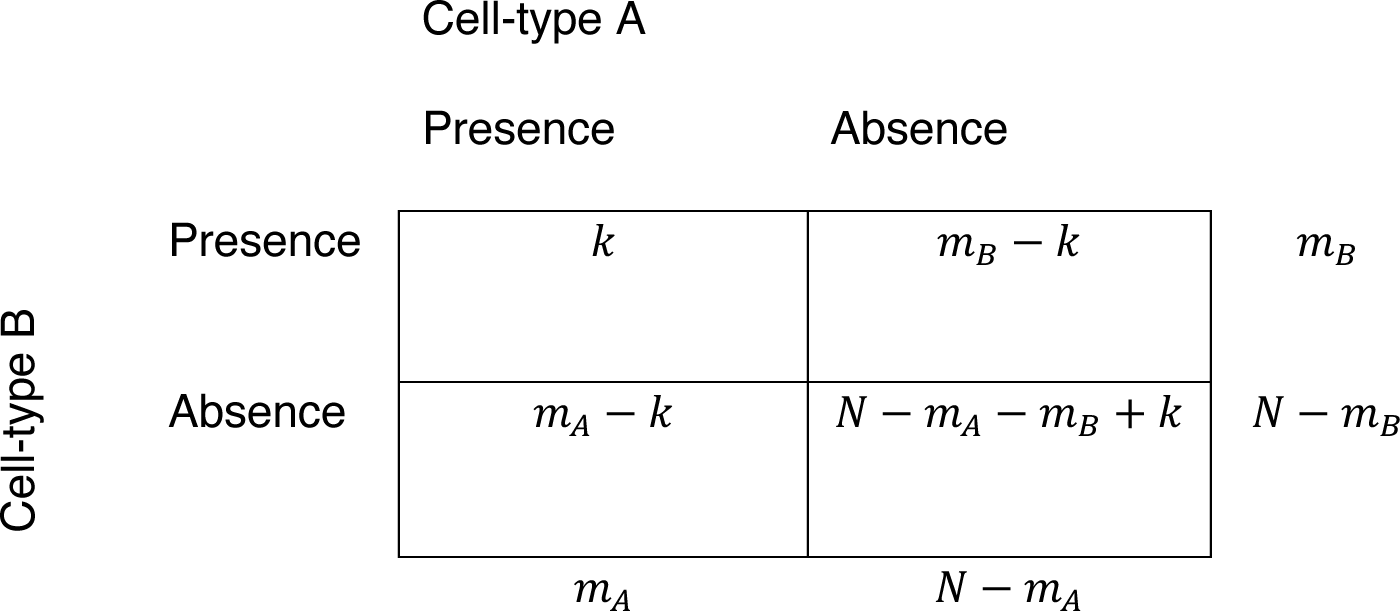

where 𝑘 denotes the number of pixels with both cell-type A and B, 𝑚_%_ denotes the number of pixels with only cell-type A, 𝑚_$_ denotes the number of pixels with only cell-type B, and 𝑁 denotes the number of all pixels. The CooccurrenceAffinity package derived that the probability of cooccurrence under non-zero affinity, 𝛼 , between cell-type A and B follows the extended hypergeometric distribution (Mainali et al., 2022; Mainali & Slud, 2022).

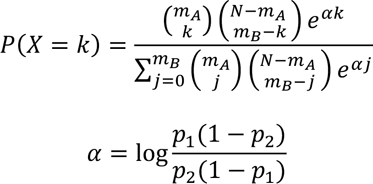

where 𝑝_1_ denotes the probability that cell-type B occupies a pixel if cell-type A is present and 𝑝_/_ denotes the probability that cell-type B occupies a pixel if cell-type A is absent.

For each contingency table, CooccurrenceAffinity computes a maximum likelihood estimate of affinity between a given cell-type pair, 𝛼̂, and test against the null hypothesis of 𝛼 = 0 (Mainali et al., 2022; Mainali & Slud, 2022). Positive 𝛼̂ indicates cooccurrence, and negative 𝛼̂ indicates separation. CooccurrenceAffinity applies an analytical null model based on hypergeometric distribution, resulting from the forementioned extended hypergeometric distribution when 𝛼 = 0, instead of permutations, allowing rapid characterization of cell-type colocalizations. P-value associated with 𝛼̂ is determined within CooccurrenceAffinity based on Blaker’s “Acceptability” function (Mainali et al., 2022). Statistically significant cooccurrence between cell-type pairs are selected based on a p-value cutoff of 0.05 or below. Confidence intervals associated with 𝛼̂ are also computed within CooccurrenceAffinity using Blaker’s method (Mainali & Slud, 2022).

Hierarchical clustering of computed 𝛼̂ values represented as a 𝑛_*j*_ x 𝑛_*j*_ symmetrical matrix, where 𝑛_̂_ is the number of categories 𝑗, was done using the hclust function in the stats package.

Runtime for SEraster alone, cell-type cooccurrence analysis alone, and combined were measured using Sys.time function in the R Base package and averaged across 5 trials. This runtime analysis was done on Mac Studio Apple M2 Ultra with 192GB unified memory (24-core CPU, 60-core GPU, 32-core Neural Engine) using 1 worker for single computer multicore parallel evaluation (BiocParallel::MulticoreParam(workers = 1)).

#### 7. Computational implementation

SEraster is implemented as an R package within the Bioconductor framework. It builds on SpatialExperiment package (Righelli et al., 2022) for processing spatial omics data within the package, Simple Feature package (Pebesma et al., 2018) for executing rasterization, Matrix package (Bates et al., n.d.) for handling sparce matrix representations, and BiocParallel package (Morgan et al., 2023) for parallelization.

#### 8. Code availability

SEraster is available as an R package on GitHub at https://github.com/JEFworks-Lab/SEraster with additional tutorials at https://JEF.works/SEraster. Code to reproduce preprocessing, analyses, and figures in this manuscript is accessible on GitHub at https://github.com/GohtaAihara/SEraster-analyses.

## Supplementary Figures

**Supplementary Figure 1.**
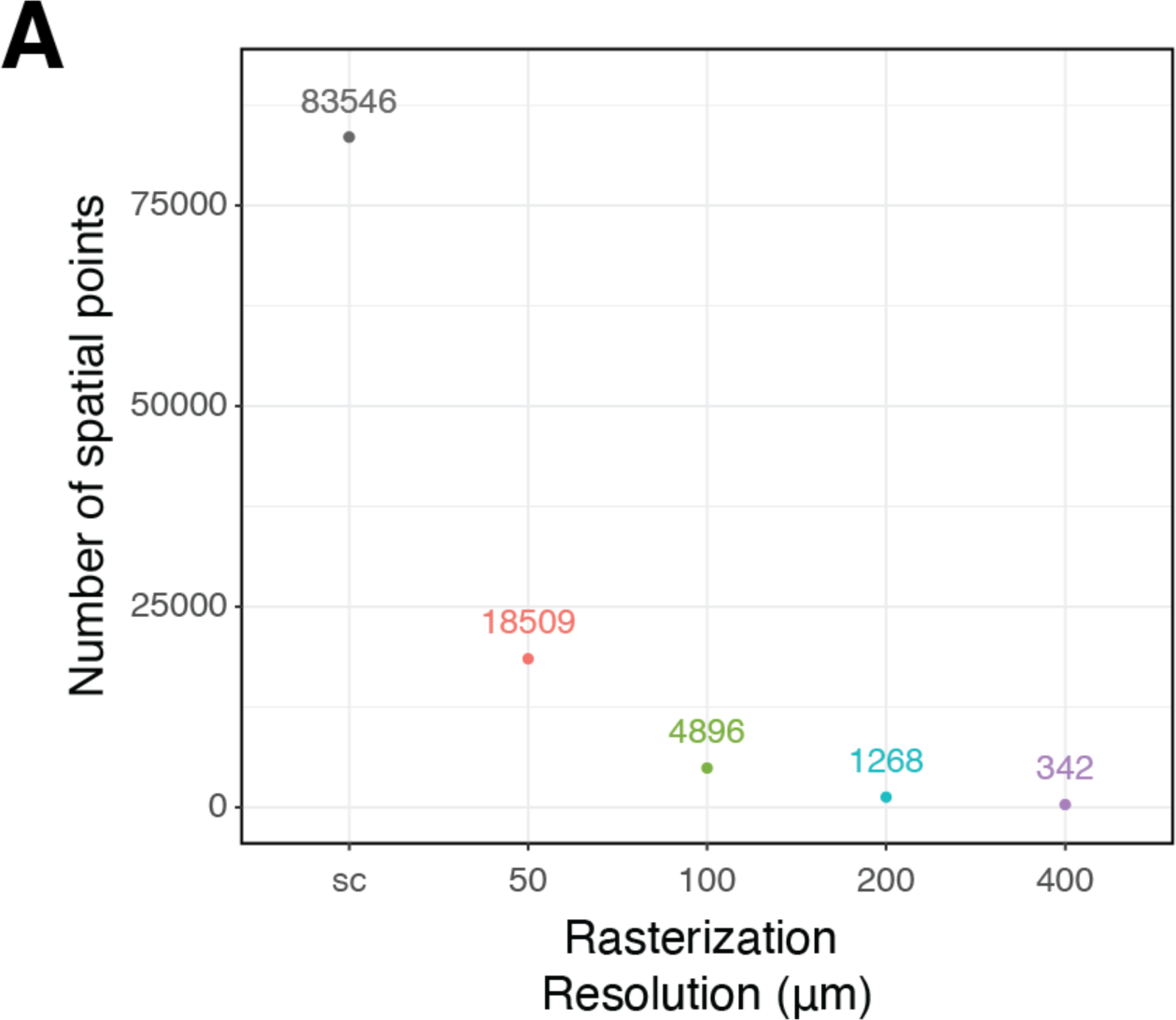
**A.** Number of spatial points for the MERFISH mouse brain dataset at single-cell resolution (sc) and rasterized resolutions (50 µm, 100 µm, 200 µm, and 400 µm). Exact numbers of spatial points are shown as text labels.

**Supplementary Figure 2.**
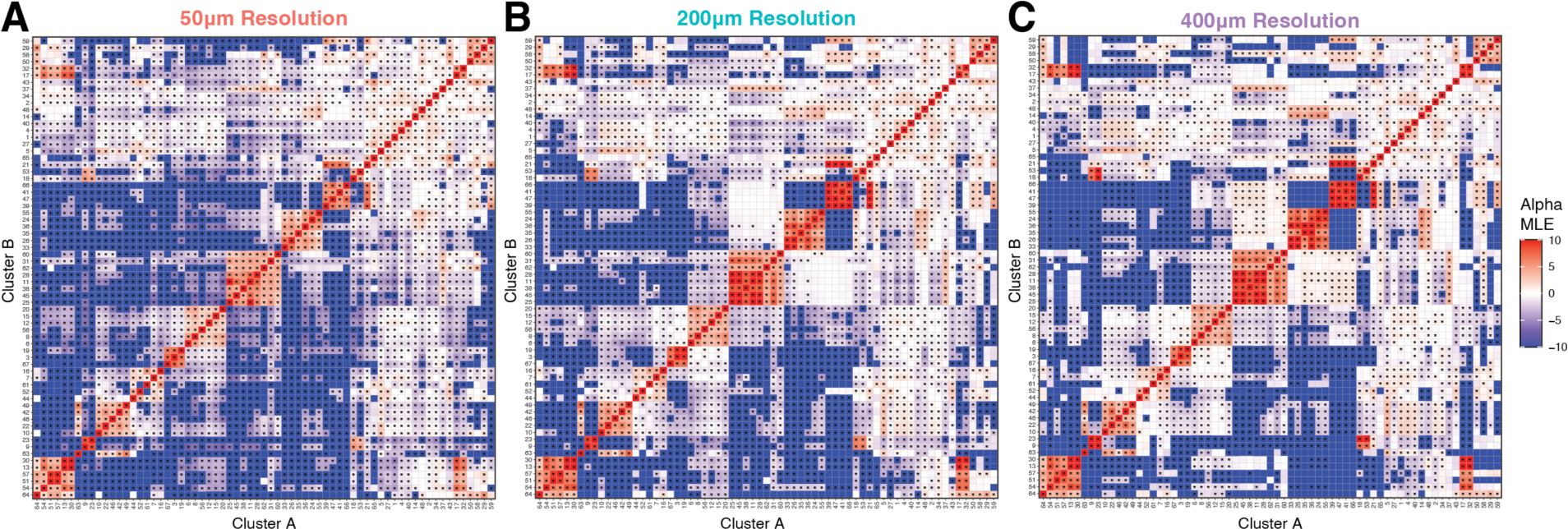
Summary of cluster cooccurrence analysis for the whole mouse pup dataset with SEraster and CooccurrenceAffinity at **A.** 50 µm, **B.** 200 µm, and **C.** 400 µm resolutions. Heatmap is colored by the maximum likelihood estimate of the affinity metric (alpha MLE) for corresponding cluster pairs. Statistically significant cooccurrences or separation (p-value ≤ 0.05) are indicated by asterisks (*).

**Supplementary Figure 3.**
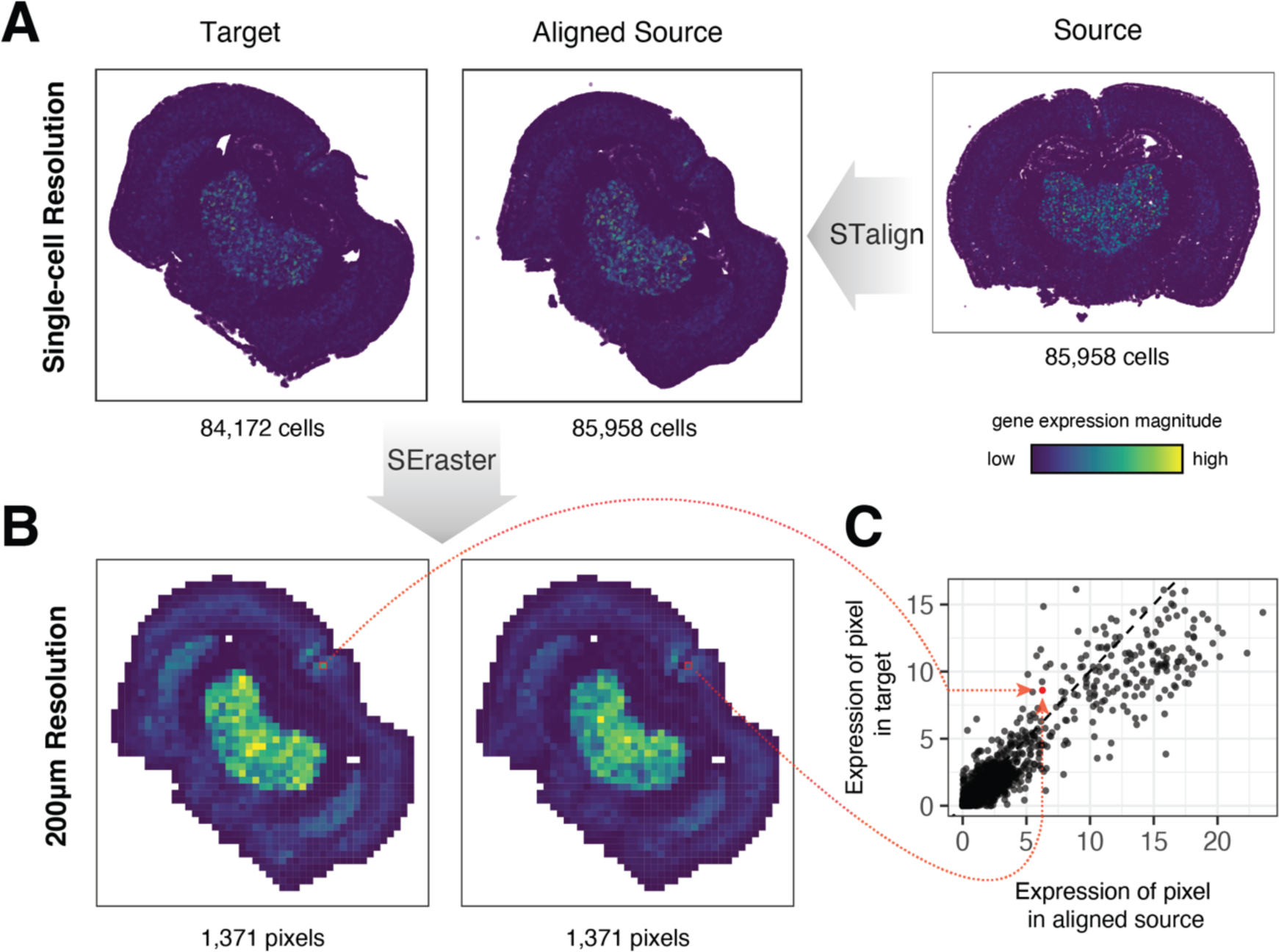
Application of SEraster to evaluate spatial correspondence of gene expression for validation of spatial alignment of single cell spatial transcriptomics datasets. A. Alignment of source (right) and target (left) single-cell resolution spatial transcriptomics datasets of coronal sections of mouse brain assayed by MERFISH using STalign resulting in an aligned source (middle). Cells colored by expression of *Grm4.* B. Joint rasterization with SEraster of target (left) and aligned source (right) datasets at 200 µm resolution. Pixels colored by rasterized expression of *Grm4.* C. Correspondence of rasterized *Grm4* counts at matched pixel locations in target and aligned source. Example of matched pixel locations indicated in red.

